# Dual RNA Polymerase I Inhibition with CX-5461 and BMH-21 Synergizes in Breast Cancer by Activating p53-Dependent Stress

**DOI:** 10.1101/2025.11.19.689351

**Authors:** Jonathan Y. Chung, Kristen N. Nguyen, Bruce A. Knutson

## Abstract

Hyperactivated ribosomal RNA (rRNA) transcription by RNA polymerase I (Pol I) is a hallmark of cancer and drives elevated ribosome biogenesis required for rapid tumor growth. Several Pol I inhibitors have been identified that induce potent anti-cancer effects. However, clinical application of the first-in-class Pol I inhibitor, CX-5461, has been limited by patient toxicity, which is comparable to other chemotherapies. Identifying synergistic drug combinations offers a promising strategy to maintain on-target anti-cancer effects while minimizing adverse reactions. Synergistic drug combinations involve drugs that enhance each other’s effect, enabling dose reduction while preserving efficacy. Synergistic drug combinations of Pol I inhibitors and other anti-cancer agents have been reported; however, it remains unclear whether Pol I inhibitors can synergize with each other. We therefore explored whether two Pol I inhibitors synergize in cancer treatment. We found that CX-5461 and BMH-21 significantly reduced MCF-7 breast cancer cell viability at clinically relevant doses. Combined treatment with these inhibitors led to profound viability defects at sub-micromolar concentrations. Our biochemical analysis showed that CX-5461 and BMH-21 combination therapy enhanced Pol I inhibition and p53 activation compared to monotherapy, promoting growth arrest and apoptosis. Collectively, our findings demonstrate that CX-5461 and BMH-21 are complementary in inhibiting Pol I, activating p53, and suppressing cancer cell growth. Based on these pre-clinical findings, dual Pol I inhibition with CX-5461 and BMH-21 represents a promising therapeutic strategy for treating cancer that is potentially both broadly applicable and tolerable.

## 2 Introduction

Hyperactivation of RNA polymerase I (Pol I) is a hallmark of cancer and contributes directly to tumor progression (Bywater et al., 2013; Zang et al., 2024). Human Pol I, a 13 subunit DNA-dependent RNA polymerase, transcribes ribosomal DNA (rDNA) loci to produce 47S ribosomal precursor ribosomal RNA (rRNA) that undergoes a series of processing and assembly steps to form mature ribosomes (Henras et al., 2015). This process, known as ribosome biogenesis, occurs primarily in the nucleolus, a specialized subnuclear condensate that serves as the site for Pol I transcription and rRNA modification. rRNA synthesis is an essential and rate-limiting step in ribosome biogenesis, and its regulation is tightly linked with cellular growth and proliferation (Scull and Schneider, 2019). In cancers, dysregulation of signaling pathways that control Pol I activity results in elevated rRNA transcription, driving increased ribosome production and protein synthesis to support rapid cell division (Bywater et al., 2013). As a result, Pol I hyperactivation has emerged as a critical driver of oncogenesis and a promising therapeutic target (Ferreira et al., 2020; Hwang and Denicourt, 2024).

Several Pol I inhibitors have been identified for use as anti-cancer therapeutics (Peltonen et al., 2010; Drygin et al., 2011; Morgado-Palacin et al., 2014; Caggiano et al., 2020). The first-in-class specific Pol I inhibitor, CX-5461, was identified through a small-molecule screen which showed that CX-5461 preferentially inhibits Pol I transcription relative to Pol II transcription (Drygin et al., 2011). Although the exact mechanism remains under investigation and may vary across cancer types, studies have shown that CX-5461 directly blocks Pol I transcription initiation (Drygin et al., 2011; Mars et al., 2020). Further studies revealed that CX-5461 induces DNA damage (Negi and Brown, 2015; Quin et al., 2016). CX-5461 intercalates DNA and stabilize G-quadruplexes (G4s), which are associated with replication fork collapse and DNA breaks (Xu et al., 2017; Liu et al., 2023). Alternatively, CX-5461 may induce DNA damage by inhibiting DNA topoisomerases, enzymes critical for resolving DNA supercoiling during replication and transcription (Cameron et al., 2024; Pan et al., 2021; Bruno et al., 2020). Despite ongoing questions about its mechanism, CX-5461 consistently exhibits strong anti-cancer activity by targeting Pol I transcription and inducing DNA damage.

In addition to CX-5461, another Pol I inhibitor, BMH-21, is currently under pre-clinical investigation for its potential use in treating cancer. Initial characterization showed that BMH-21 intercalates into GC-rich DNA, inhibits Pol I transcription, and activates a transcription elongation checkpoint that induces proteasomal degradation of the Pol I catalytic subunit, RPA194 (Peltonen et al., 2010, 2014; Pitts et al., 2022). Further characterization demonstrated that BMH-21 reduces both initiation and elongation of rRNA transcription and promotes polymerase pausing (Jacobs et al., 2022). In contrast to CX-5461, BMH-21 does not appear to induce DNA breaks or activate an ATM-dependent DNA damage response (Colis et al., 2014). However, BMH-21 has been shown to stabilize G4s (Musso et al., 2018; Mazzini et al., 2019) and modulate topoisomerase II activity and localization (Morotomi-Yano and Yano, 2021; Espinoza et al., 2024). Through its multi-faceted disruption of Pol I transcription and DNA topology, BMH-21 represents a promising and mechanistically distinct approach to targeting Pol I in cancer therapy.

While CX-5461 and BMH-21 differ in their mechanisms, both activate p53-dependent tumor suppression. Inhibition of Pol I by either CX-5461 or BMH-21 activates of the major tumor suppressor p53 via the nucleolar surveillance pathway (NSP) wherein impaired rRNA synthesis leads to inhibition of MDM2, a key negative regulator of p53 protein stability (Deisenroth and Zhang, 2010; Bywater et al., 2012). Stabilized p53 then initiates a tumor-suppressive transcriptional program that includes genes involved in genome maintenance, growth arrest, and apoptosis. In addition, CX-5461 activates p53 through a second mechanism by inducing DNA damage and engaging the ATM-dependent DNA damage response (Cheng and Chen, 2010; Sanij et al., 2020). These complementary mechanisms raise the possibility that combining CX-5461 and BMH-21 could enhance p53 activation and synergistically promote tumor suppression.

Synergistic drug combinations enhance therapeutic efficacy by producing effects greater than the sum of each drug alone. These combinations allow for lower individual drug doses, which can reduce off-target toxicity, improve specificity, and prevent chemoresistance (Lehàr et al., 2009; Mokhtari et al., 2017). CX-5461, currently in clinical trials, has shown dose-limiting side effects such as nausea and phototoxicity (Hilton et al., 2022). Identifying synergistic partners for CX-5461 could mitigate these toxicities while maintaining anti-cancer potency by reducing the dose of CX-5461 necessary to elicit a therapeutic effect. Although CX-5461 and BMH-21 both target Pol I and show promise individually, their combined therapeutic potential remains unexplored. Clinically, combining agents that act on the same pathway has proven effective. Trastuzumab and pertuzumab target different domains of the HER2 receptor and form a standard treatment for HER2-positive breast cancer (Jagosky and Tan, 2021), while BRAF and MEK inhibitors are co-administered to treat BRAF-mutated metastatic melanoma, improving outcomes and reducing toxicity (Eroglu and Ribas, 2016). Dual targeting of Pol I through distinct mechanisms may offer a powerful therapeutic strategy to enhance efficacy and limit adverse effects.

In this study, we investigated whether co-targeting RNA polymerase I with CX-5461 and BMH-21 offers a synergistic strategy for cancer therapy. Given that these inhibitors act through distinct mechanisms, yet both activate p53-dependent tumor suppression, we hypothesized that their combination could maintain efficacy with lower doses. Our results demonstrated that CX-5461 and BMH-21 synergize at therapeutic concentrations to significantly reduce breast cancer cell viability. This combination enhanced Pol I inhibition, increased p53 stabilization, and promoted robust growth arrest and apoptosis. Importantly, comparable reductions in cancer cell viability were achieved with substantially lower doses of each drug when used together. These findings support the use of combined Pol I inhibition with CX-5461 and BMH-21 as a potent and potentially more tolerable approach to tumor suppression.

## 3 Materials and Methods

### 3.1 Cell Culture

MCF-7 cells were obtained from ATCC (Manassas, VA). MCF-7 cells were cultured in Dulbecco’s Modified Eagle’s Medium – High Glucose (D5796, Sigma-Aldrich, Darmstadt, Germany) supplemented with 10% fetal bovine serum (F2442, Sigma-Aldrich). Cells were cultured at 37°C in a 5% CO_2_ humidified atmosphere. Cells were checked for mycoplasma contamination by PCR as described by Siegl et al. (Siegl et al., 2023).

### 3.2 Viability Assays and Synergy Determination

CX-5461 (HY-13323) and BMH-21 (HY-12484) were purchased from MedChemExpress (Monmouth Junction, NJ). CX-5461 was dissolved in 50 mM sodium phosphate (pH 3.5). BMH-21 was dissolved in DMSO (34869, Sigma-Aldrich).

Initial single-agent viability assays were performed in 24-well plates (229124, CellTreat, Pepperell, MA). 25,000 cells were seeded 16 hours prior to drug treatment of 3 replicates. Combination treatment viability assays were performed in 96-well plates (229197, CellTreat). 5,000 cells were seeded 16 hours prior to drug treatment of 4 replicates. Cell viability was assessed by MTT assay. MTT (475989, Sigma-Aldrich) was dissolved in Dulbecco’s Phosphate Buffered Saline (DPBS) (D8537, Sigma-Aldrich). Following drug treatment for 48 hours, cells were treated with 0.5 mg/mL MTT for 2 hours in growth conditions. MTT formazan crystals were solubilized in DMSO. MTT formazan absorbance was measured at 562 nm on a SpectraMax® i3x microplate reader (Molecular Devices, San Jose, CA). Viability measurements from combination treatment assays were normalized to the vehicle-treated control and analyzed using the SynergyFinder Plus platform (Zheng et al., 2022).

### 3.3 RNA Metabolic Labeling

RNA metabolic labeling was adapted from McNamar et al. (McNamar et al., 2023) and performed in 8-well chamber slides (80841, ibidi, Gräfelfing, Germany). Slides were coated with 0.01% poly-L-lysine (P4707, Sigma-Aldrich). 50,000 cells were seeded per well 16 hours prior to drug treatment. Following treatment, cells were treated for 1 hour with 1 mM 5-ethynyl uridine (EU) (CCT-1261, Vector Labs, Newark, CA) in growth conditions. Cells were fixed with 4% formaldehyde in PBS (FB002, Invitrogen) for 15 minutes at room temperature. Cells were permeabilized and blocked with 5% bovine serum albumin (BSA) (BP1600-100, Fisher Scientific) and 0.5% Triton X-100 (X100, Sigma-Aldrich) in PBS for 20 minutes at room temperature. Cells were washed with PBS-BSA (3% BSA in PBS) for 5 minutes. Click reaction solution comprised 4 mM copper sulfate (1.02790, Sigma-Aldrich), 40 mM sodium-L-ascorbate (11140, Sigma-Aldrich), and 5 μM AZDye 594 azide (CCT-1283, Vector Labs) in PBS. Cells were incubated in the dark with click reaction solution for 1 hour at room temperature. Cells were washed with PBS-BSA for 5 minutes prior to counterstaining with 2μg/mL Hoescht 33342 (H3570, Thermo Fisher Scientific, Waltham, MA) in PBS-BSA for 15 minutes at room temperature. Cells were washed once with PBS-BSA and 3 times with ddH_2_O for 5 minutes. Cells were mounted with VECTASHIELD® PLUS antifade mounting medium (H-1900, Vector Labs). Slides were imaged with a Leica TCS SP8 confocal microscope at 20x magnification.

### 3.4 Western Blotting

For Western blots, 500,000 cells were seeded in 6-well plates (229105, CellTreat) 16 hours prior to drug treatment. Following treatment, cells were lysed with RIPA buffer (DeCaprio and Kohl, 2021) with 5 mM EDTA, protease inhibitors, and phosphatase inhibitors (B15001, Selleck Chemicals LLC, Houston, TX). Lysates were run on SDS-PAGE gels and transferred to PVDF membranes. Total protein was detected using Revert™ 520 total protein stain (926-10011, LI-COR, Lincoln, NE). Primary antibodies used in Western blots assays were diluted in TBST and incubated with membranes overnight at 4°C. Primary antibodies included anti-RPA194 (1:1,000 dilution) (sc-48385, Santa Cruz Biotechnology (SCBT), Dallas, TX), anti-p53 (1:2,000 dilution) (sc-126, SCBT), anti-p21 (1:200 dilution) (sc-6246, SCBT), anti-caspase-7 (1:500 dilution) (sc-56063, SCBT), p-histone H2A.X (1:200 dilution) (sc-517348, SCBT). Blots were incubated 2 hours at room temperature with IRDye® 680RD goat anti-mouse IgG secondary antibody diluted 1:5,000 in TBST (926-68070, LI-COR) and imaged using the Odyssey M imager (LI-COR). Densitometry was performed using ImageStudio (LI-COR). Relative abundance of proteins were compared across treatment conditions by normalizing to total protein abundance.

### 3.5 Statistical Analysis

For viability assays, control cell viability was compared to treated cell viability using RM one-way ANOVA with Dunnett’s multiple comparisons test. One-sample two-tailed t-tests were applied to compare observed viability to the predicted viability of each synergy model. For EU incorporation assays, nuclear mean fluorescence intensities of single cells were compared across treatments using Brown-Forsythe and Welch ANOVA tests and Dunnett’s T3 multiple comparisons test. For Western blotting assays, relative target protein abundances were compared across treatments RM one-way ANOVA with Dunnett’s multiple comparisons test. All statistical analyses was performed with GraphPad Prism (version 10.4.1) (GraphPad, Boston, MA).

## 4 Results

### 4.1 Combined Pol I inhibition with CX-5461 and BMH-21 reduces MCF-7 cell viability

To investigate the therapeutic potential of dual Pol I inhibition, we examined the effects of combining CX-5461 and BMH-21 on MCF-7 luminal A breast cancer (BC) cell viability **(Figure 1)**. Luminal A BCs constitute most BC diagnoses, and despite their initial favorable response to hormone therapy and resection, up to 10% of luminal A BCs metastasize within 10 years of their diagnosis (van Maaren et al., 2019). Homozygous p53 wild-type MCF-7 cells were selected for this study as ∼80% of luminal BCs are p53 wild-type and Pol I inhibitors activate a p53-dependent nucleolar surveillance pathway (Mueller et al., 2023; Bywater et al., 2012). Up to 80% of luminal BCs that do not respond to preoperative hormone therapy are p53 wild-type (Mueller et al., 2023). Frequency of metastasis and acquired resistance to hormone therapies underscores a need for effective and generalizable chemotherapies for luminal BCs (Clarke et al., 2015).

**Figure 1.**
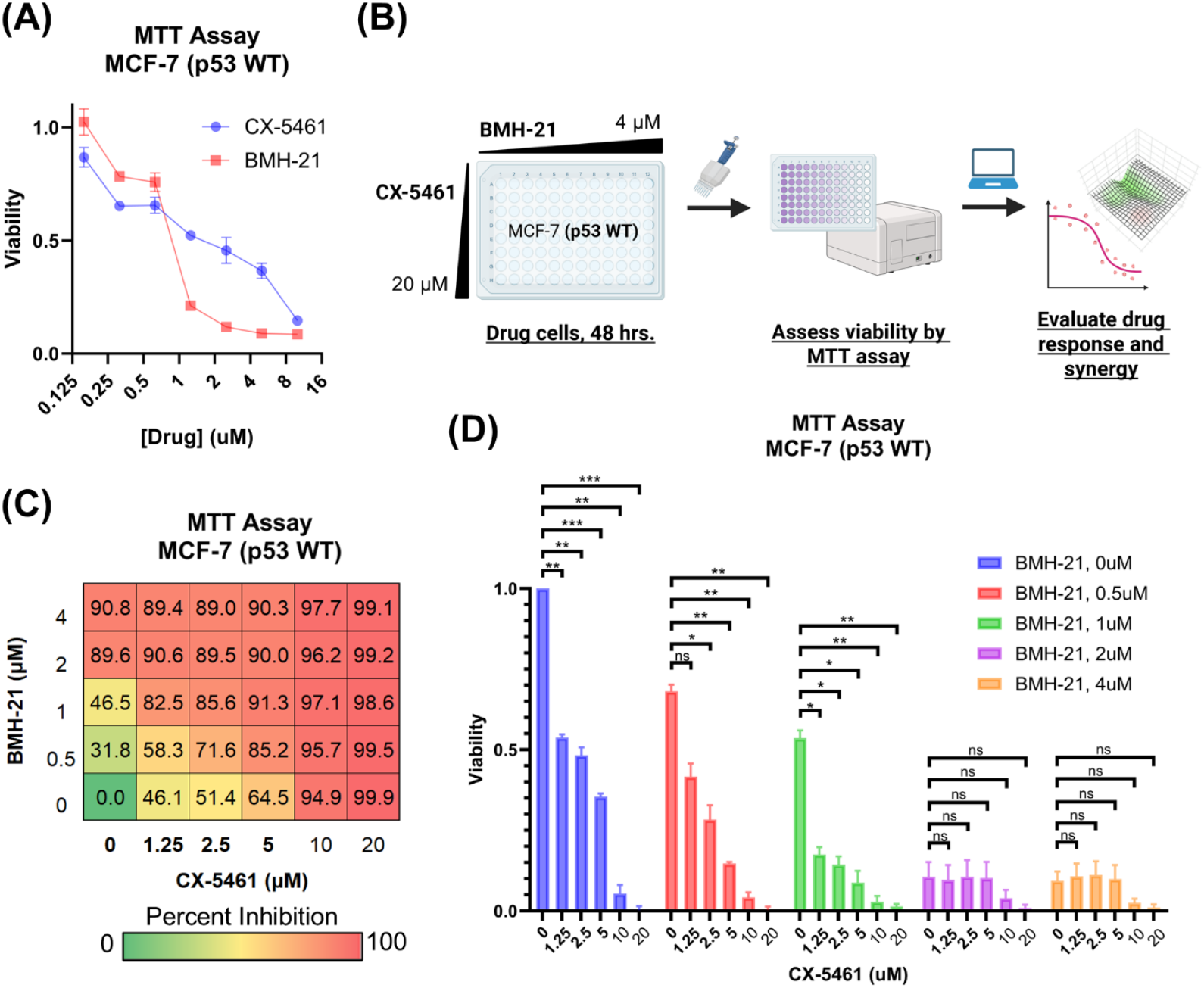
CX-5461 and BMH-21 combinations reduce MCF-7 cell viability. **(A)** Viability of MCF-7 luminal breast cancer cells treated with increasing doses of CX-5461 or BMH-21. Viability was measured by MTT assay following treatment of cells for 48 hours. Error bars represent the standard deviation across three technical replicates. **(B)** CX-5461 and BMH-21 synergy evaluation workflow. MCF-7 cells were treated with combinations of BMH-21 (0 – 4 μM) and CX-5461 (0 – 20 μM) for 48 hours prior to viability measurement. Viability measurements were evaluated using SynergyFinder Plus to determine synergy scores. **(C)** Dose-response matrix indicating percent inhibition of MCF-7 cells, normalized to vehicle-treated cells, following 48-hour treatment with CX-5461 and BMH-21. Serum concentrations of CX-5461 achievable in clinical trials are bolded. Values represent the average of three biological replicates composed of four technical replicates each. **(D)** Viability of MCF-7 cells treated as in (C). Error bars represent the standard error of the mean. Viability was compared using RM one-way ANOVA with Dunnett’s multiple comparisons test, with p-value < 0.05 (*), p-value < 0.01 (**), p-value < 0.001 (***), and not significant (n.s.) indicated.

We first evaluated the cytotoxicity of each drug individually and found that 48-hour treatment with BMH-21 induced significant cell death at 2 μM, whereas CX-5461 required concentrations up to 10 μM to achieve a comparable effect (**Figure 1A**). Based on these responses, we performed a checkerboard synergy assay using dose ranges that captured the full activity spectrum of each drug (**Figure 1B**). Co-treatment with CX-5461 and BMH-21 led to a greater reduction in MCF-7 cell viability compared to either agent alone (**Figure 1C**), indicating a cooperative interaction between the two inhibitors. While 20 μM CX-5461 alone cleared nearly all cells, addition of BMH-21 to lower concentrations of CX-5461 produced comparable or greater reductions in viability than CX-5461 alone, underscoring the enhanced efficacy of dual Pol I inhibition. Although a small fraction of cells remained following BMH-21 treatment alone (2 μM and 4 μM), these cells remained sensitive to higher doses of CX-5461, further supporting the benefit of combined therapy. These findings demonstrate that combining CX-5461 and BMH-21 promotes robust suppression of breast cancer cell viability, supporting their potential use in combination-based cancer therapy.

### 4.2 CX-5461 and BMH-21 exhibit dose-dependent synergy in MCF-7 cells

To assess the nature of the interaction between CX-5461 and BMH-21, we analyzed MCF-7 cell viability data using SynergyFinder Plus, a platform that supports multiple reference models for drug interaction analysis (**Figure 2**) (Zheng et al., 2022). Given the lack of consensus on the most appropriate synergy model, we employed all four available models in SynergyFinder Plus (Loewe additivity, Bliss independence, zero-interaction potency (ZIP), and highest single agent (HSA)) to ensure a comprehensive evaluation (Tang et al., 2015; Yadav et al., 2015). Mean synergy scores were calculated across three independent experiments for each dose combination (**Figure 2A**). Across the full dose range, average synergy scores were close to zero for the Loewe, Bliss, and ZIP models, indicating largely additive effects. The HSA model, which compares combination efficacy to the most effective single agent, yielded scores greater than zero, suggesting enhancement in viability reduction when the drugs were combined. These results indicate that CX-5461 and BMH-21 exhibit additive to mildly synergistic interactions overall, with potential for synergy at select doses.

**Figure 2.**
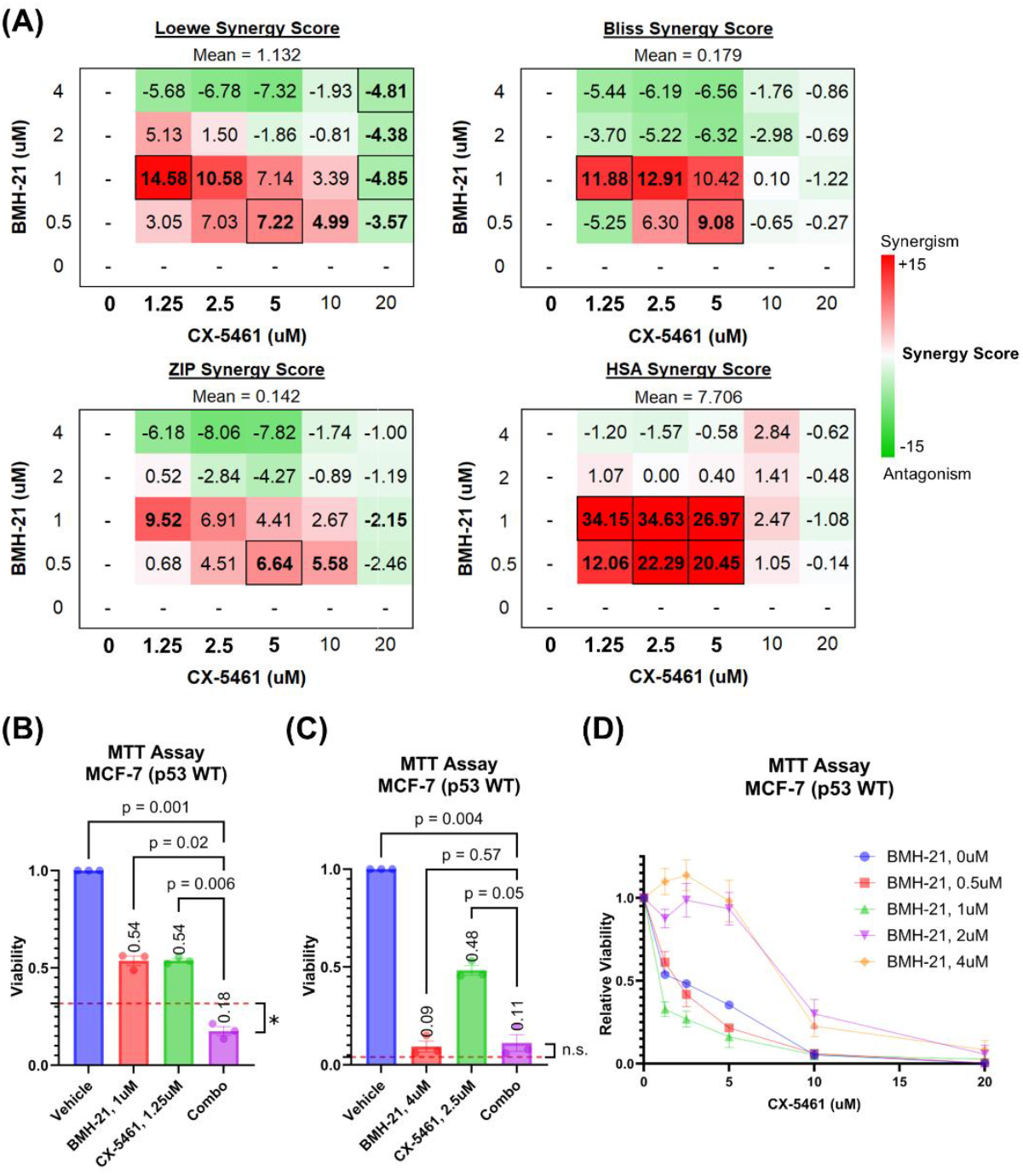
CX-5461 and BMH-21 synergize at low to moderate doses. Reported values and points represent the average of three biological replicates composed of four technical replicates each. Error bars represent the standard error of the mean. **(A)** Synergy scores of all CX-5461 and BMH-21 dose combinations. Loewe additivity, Bliss independence, zero-interaction potency (ZIP), and highest single-agent (HSA) reference scores were calculated using SynergyFinder Plus to evaluate synergy. Synergy scores represent the difference between the observed viability and predicted viability estimated by each reference model, with positive (red) values representing enhanced viability reduction and synergy. Serum concentrations of CX-5461 achievable in clinical trials are bolded. Synergy scores which significantly deviated (p < 0.05) are outlined and scores which nearly met significance (p < 0.1) are bolded. Significance was determined using one-sample two-tailed t-test. **(B)** Viability of MCF-7 cells treated with the most synergistic doses of CX-5461 and BMH-21. The red dashed line represents the viability of combined treatment predicted by the Loewe additivity reference model. Cell viabilities were compared using RM one-way ANOVA with Dunnett’s multiple comparisons test. Cell viability following combination therapy was compared to the predicted viability using one-sample two-tailed t-test, with p-value < 0.05 (*) and not significant (n.s.) indicated. **(C)** Viability of MCF-7 cells treated with the most antagonistic doses of CX-5461 and BMH-21, as in (B). **(D)** CX-5461 dose-viability relationship of cells treated with varying doses of BMH-21. Cell viabilities were normalized to the viability of cells receiving the same dose of BMH-21 without CX-5461.

To explore how drug concentration influences synergy, we examined synergy scores for individual dose combinations of CX-5461 and BMH-21. Notably, the two drugs synergized at low to moderate concentrations but exhibited antagonism at higher doses, resulting in an overall average synergy score near zero across all combinations (**Figure 2A**). Synergistic interactions were most pronounced within a range of 0.5–1 μM for BMH-21 and 1.25–10 μM for CX-5461. These ranges include the drugs’ respective single-agent IC50 values (0.84 μM for BMH-21 and 1.47 μM for CX-5461) and align with serum concentrations of CX-5461 reported in phase I clinical trials (Khot et al., 2019; Hilton et al., 2022).

The most potent synergistic combination (1.25 μM CX-5461 with 1 μM BMH-21) reduced cell viability by 82%, significantly surpassing the effect predicted by the Loewe additivity model (p = 0.025; **Figure 2B**). In contrast, combinations exceeding 1 μM BMH-21 and 10 μM CX-5461 were generally antagonistic. For example, 2.5 μM CX-5461 with 4 μM BMH-21 reduced viability by 89%, a level comparable to BMH-21 monotherapy and 5–6% below Loewe model predictions (p = 0.236; **Figure 2C**). Although these high-dose combinations were classified as antagonistic by synergy models, they still markedly suppressed cancer cell viability. Importantly, the antagonistic dose range for CX-5461 (10–20 μM) exceeds the upper limits of clinically achievable serum concentrations (Hilton et al., 2022), reinforcing the translational relevance of the lower-dose synergistic combinations.

Next, we assessed how BMH-21 influences the potency of CX-5461. Dose-response curves for CX-5461, normalized to cells not treated with CX-5461, showed a clear leftward shift in the presence of BMH-21 up to 1 μM, indicating enhanced CX-5461 potency (**Figure 2D**). Curve fitting confirmed that increasing BMH-21 concentrations up to 1 μM lowered the CX-5461 IC50 **(Supplementary Table 1**). However, at 2 μM and 4 μM BMH-21, the CX-5461 IC50 increased, suggesting that higher doses of CX-5461 were required to eliminate the small number of cells that remained after high-dose BMH-21 treatment. These residual cells, though few (**Figure 1C, 2D**), were effectively cleared with elevated CX-5461 concentrations. Together, these findings demonstrate that BMH-21 enhances CX-5461 potency at low to moderate doses, driving modest synergy, whereas high-dose combinations display antagonistic interactions despite maintaining robust overall cytotoxicity.

### 4.3 CX-5461 and BMH-21 combination treatment suppresses nascent RNA synthesis

To better understand how the drug combination disrupts cancer cell function, we examined its impact on RNA synthesis. Specifically, we measured nascent RNA production by using ethynyluridine (EU) incorporation detected by click chemistry (Jao and Salic, 2008). Although EU is incorporated into all newly transcribed RNAs, it primarily reflects rRNA synthesis by RNA polymerase I (Pol I), which accounts for the majority of total transcription in the cell (Moss and Stefanovsky, 2002). As expected, EU uptake was limited to cell nuclear compartments. A 3-hour treatment with 0.5 μM CX-5461 and 0.5 μM BMH-21 led to a complete loss of EU signal in nucleoli, in contrast to the strong nucleolar labeling seen in vehicle-treated cells (**Figure 3A**). Quantification of nuclear EU fluorescence confirmed that the combination significantly reduced RNA synthesis compared to either drug alone or vehicle control (**Figure 3B**). These results indicate that dual Pol I inhibition more effectively disrupts rRNA transcription than monotherapy, providing further evidence of the synergy between CX-5461 and BMH-21.

**Figure 3.**
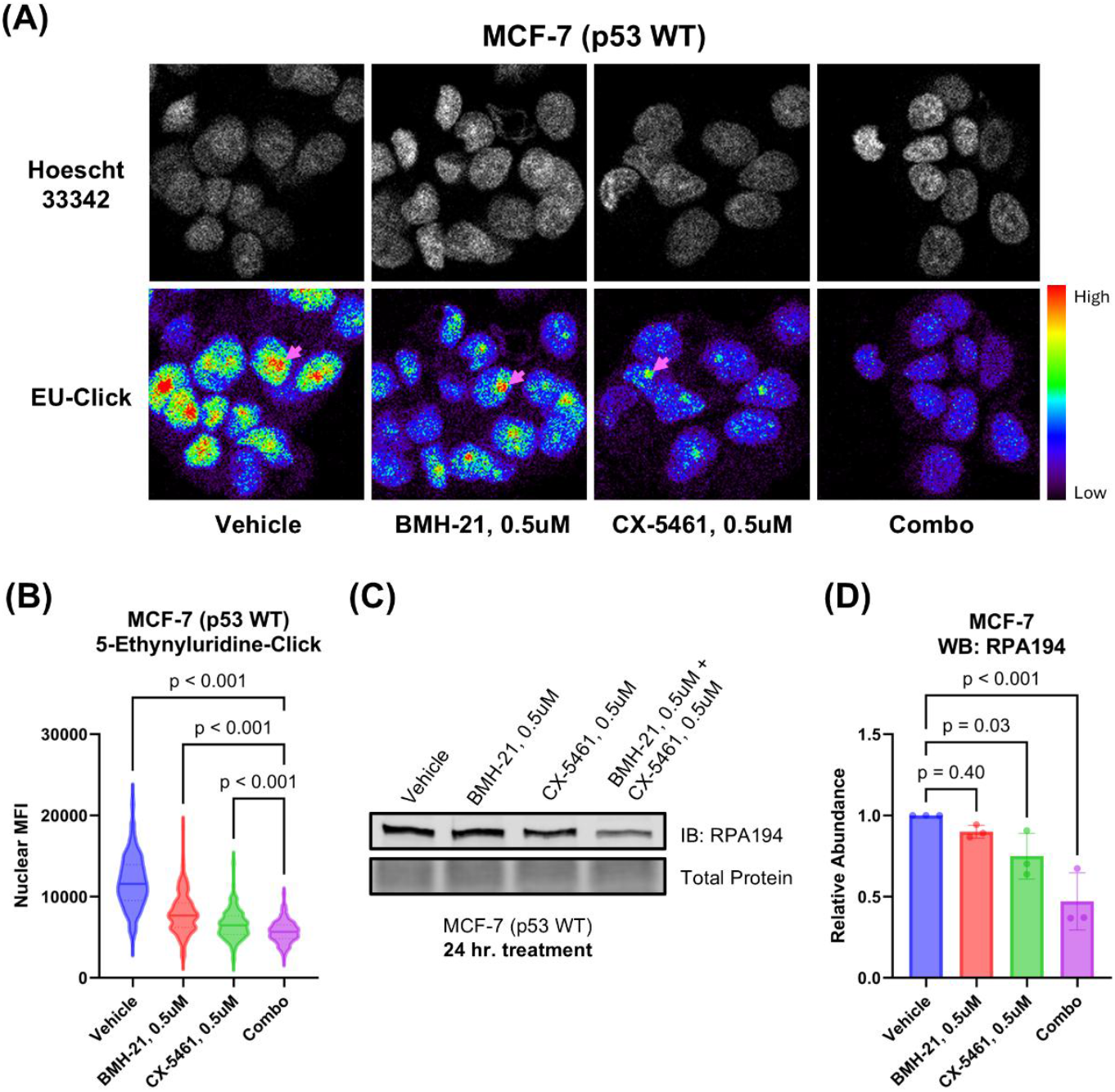
CX-5461 and BMH-21 synergistic combinations enhance Pol I inhibition. **(A)** Ethynyl uridine (EU) incorporation assay measuring nascent RNA synthesis. MCF-7 cells were treated for 3 hours prior to addition of EU. Examples of nucleoli are indicated with pink arrows. Representative images are shown. **(B)** Quantitation of nuclear EU signal in (A). Nuclear mean fluorescence intensity (MFI) of single cells were compared across treatments using Brown-Forsythe and Welch ANOVA tests and Dunnett’s T3 multiple comparisons test. At least 100 cells were analyzed for each treatment condition. **(C)** Western blot probing for the large catalytic subunit of Pol I (RPA194) as a measure of Pol I inhibition. MCF-7 cells were treated 24 hours prior to harvesting. A representative Western blot is shown. **(D)** Densitometry of (C). Bars represent the average of three biological replicates and error bars represent the standard deviation. Relative abundance was compared using RM one-way ANOVA with Dunnett’s multiple comparisons test.

### 4.4 CX-5461 and BMH-21 combination treatment reduces Pol I abundance

To investigate how the drug combination affects Pol I stability, we examined levels of the catalytic subunit of Pol I, RPA194, by Western blot (**Figure 3C–D**). BMH-21 is known to promote proteasomal degradation of RPA194 through activation of a transcription elongation checkpoint and FBXL14-mediated ubiquitination (Pitts et al., 2022). CX-5461 may also induce RPA194 degradation, though less efficiently (Pitts et al., 2022). In addition, RPA194 levels can be downregulated transcriptionally via nucleolar stress-induced repression of c-MYC, a key regulator of ribosome biogenesis (Liao et al., 2014). Treatment with 0.5 μM CX-5461 and 0.5 μM BMH-21 for 24 hours significantly reduced RPA194 abundance (p < 0.001) by approximately twofold, whereas monotherapies resulted in only marginal decreases (**Fig. 3D**). These findings suggest that the combination of CX-5461 and BMH-21 more effectively downregulates Pol I than either agent alone.

### 4.5 CX-5461 and BMH-21 cooperatively enhances DNA damage signaling

As CX-5461 and BMH-21 are DNA intercalators that have been implicated in causing DNA and chromatin damage, we evaluated activation of the DNA damage response (DDR) by combination treatment with CX-5461 and BMH-21 (Colis et al., 2014; Espinoza et al., 2024). To evaluate activation of DDR, we examined treated cells for the presence of phosphorylated H2A histone family member X (H2A.X) by Western blot. Phosphorylation of H2A.X at serine 139, referred to as γH2A.X, serves as an early indicator of DDR which coordinates recruitment of DNA damage repair machinery (Collins et al., 2020). Interestingly, treating MCF-7 cells with both 1 μM CX-5461 and 1 μM BMH-21 for 48 hours enhanced γH2A.X abundance compared to the monotherapies (p = 0.01, **Supplementary Figure 1**). Therefore, combined treatment with CX-5461 and BMH-21 may enhance DDR activation compared to monotherapies. Indeed, the nucleolus, Pol I transcription, and DNA damage along with its repair are intimately related and synergy between CX-5461 and BMH-21 could be mediated by crosstalk between Pol I and DNA damage pathways (Lindström et al., 2018).

### 4.6 Pol I inhibitor combination treatment induces p53-dependent growth arrest and apoptosis

Because both the nucleolar surveillance pathway (NSP) and DNA damage response (DDR) converge on p53 activation (Zhang et al., 2003; Cheng and Chen, 2010), we assessed whether CX-5461 and BMH-21 enhance p53 signaling (**Figure 4**). In the NSP, reduced rRNA synthesis leads to MDM2 inhibition and subsequent p53 stabilization, which in turn drives transcription of p53 target genes such as the cell cycle inhibitor p21 (El-Deiry et al., 1994). We therefore measured the abundance of p53 and p21 to assess p53 activation in combination-treated MCF-7 cells. Treatment with 0.5 μM CX-5461 and 0.5 μM BMH-21 for 24 hours significantly increased p53 (p = 0.02) and p21 (p < 0.001) protein levels compared to monotherapies (**Figure 4A–C**), suggesting enhanced p53 activation at synergistic doses. These results imply that the observed reduction in cell viability is, at least in part, driven by p53-dependent p21 upregulation and G1 phase arrest (Waldman et al., 1995).

**Figure 4.**
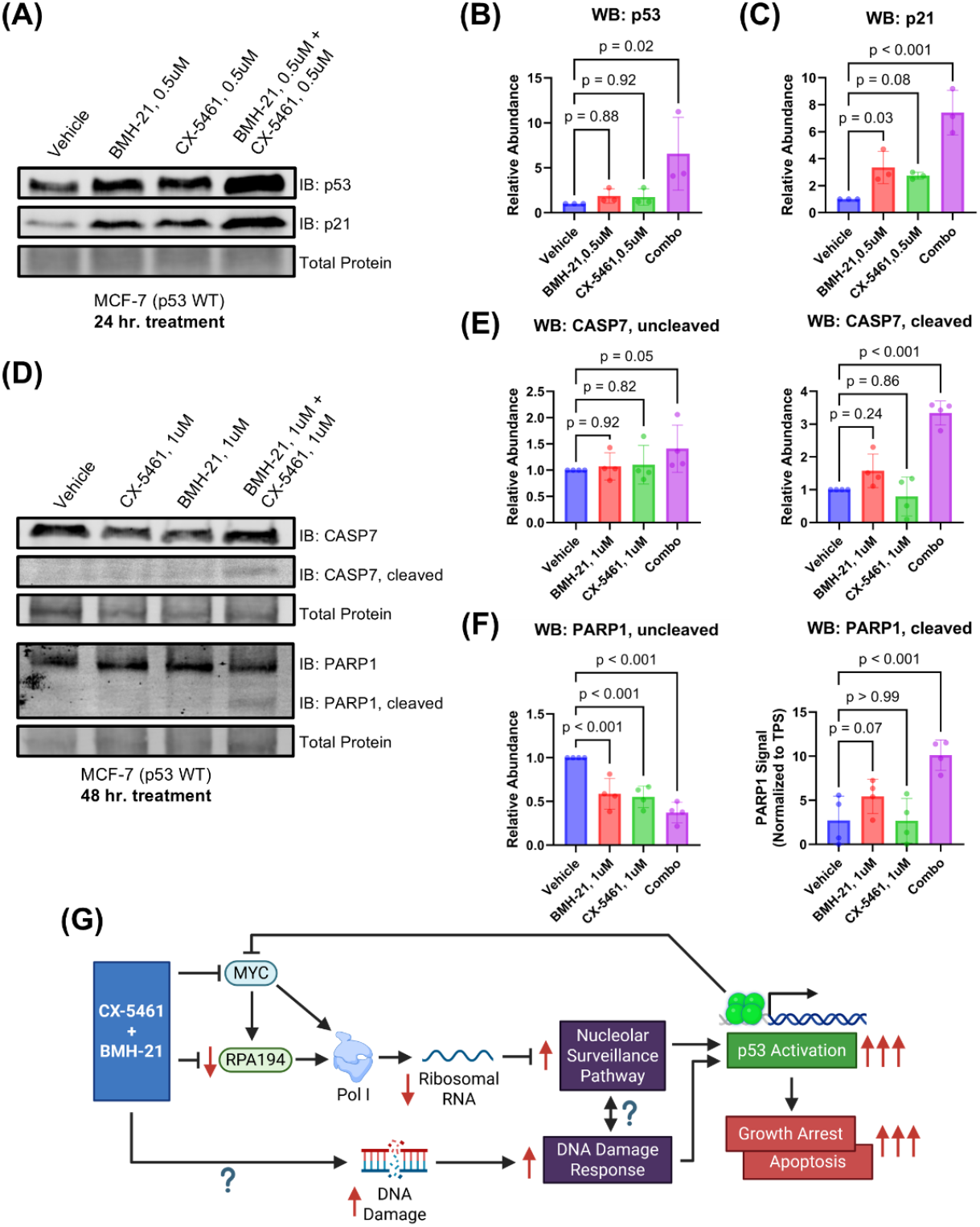
CX-5461 and BMH-21 synergistic combinations enhance p53-dependent growth suppression and apoptosis. **(A)** Western blot probing for p53 and p21 abundance as measures of p53 activation. MCF-7 cells were treated for 24 hours prior to harvesting. A representative Western blot is shown. **(B, C)** Densitometry of (A). Bars represent the average of three biological replicates and error bars represent the standard deviation. Relative abundance was compared using RM one-way ANOVA with Dunnett’s multiple comparisons test. **(D)** Western blot probing for caspase-7 (CASP7) and poly(ADP-ribose) polymerase 1 (PARP1) total and cleaved forms as a measure of apoptosis induction. MCF-7 cells were treated for 48 hours with 1 μM CX-5461 and/or 1 μM BMH-21 prior to harvesting. Representative Western blots are shown. **(E, F)** Densitometry of (D). Bars represent the average of four biological replicates and error bars represent the standard deviation. Relative abundance was compared using RM one-way ANOVA with Dunnett’s multiple comparisons test. **(G)** Model of CX-5461 and BMH-21 combination therapy mechanism of action. Created in BioRender. Chung, J. (2025) https://BioRender.com/jc65dsz

We next examined whether combination treatment also activates intrinsic apoptosis, a process initiated by sustained p53 activation (Chen, 2016). To evaluate apoptosis, we measured cleavage of caspase-7 and its downstream substrate PARP1, both hallmarks of apoptotic execution (Germain et al., 1999; Lamkanfi and Kanneganti, 2010; Chung and Knutson, 2025). At 0.5 μM dosing, no cleaved caspase-7 or PARP1 was detected at 24- and 48-hour timepoints, indicating that lower doses primarily trigger growth arrest **(Supplementary Figure 2**). Treatment with 1 μM CX-5461 and 1 μM BMH-21 for 48 hours induced substantial cleavage of both caspase-7 and PARP1, with levels exceeding those observed in monotherapy conditions (p < 0.001; **Figure 4D–F**). Un-cleaved caspase-7 was slightly increased with combination treatment (p = 0.05), while un-cleaved PARP1 was reduced, consistent with cleavage and inactivation (p < 0.001). Together, these results demonstrate that CX-5461 and BMH-21 cooperatively activate p53 signaling to induce both cell cycle arrest and apoptosis. These dual outcomes are consistent with canonical p53 tumor suppressor functions and likely underlie the synergistic reduction in cancer cell viability observed with this combination.

## 5 Discussion

In this study, we demonstrated that the first-in-class RNA polymerase I inhibitors CX-5461 and BMH-21 worked synergistically to reduce the viability of p53 wild-type MCF-7 breast cancer cells. The most pronounced synergy occurred at low to moderate doses that were close to the IC50 values for each inhibitor. We also noted antagonism between CX-5461 and BMH-21 at higher doses of CX-5461 (over 10 μM) which are not clinically achievable. Importantly, concentrations of CX-5461 we found to modestly synergize with BMH-21 are clinically achievable, as demonstrated in phase I human clinical trials (Khot et al., 2019; Hilton et al., 2022). At these doses, the combination treatment significantly enhanced p53 activation, which led to both growth suppression and apoptosis. These findings suggest that the synergy we observed in our viability assays is driven by enhanced activation of p53-dependent stress pathways. We suspect that antagonism occurs with doses of CX-5461 and BMH-21 that induce maximal p53 activation that is not further enhanced with addition of the other Pol I inhibitor.

Our biochemical analyses indicate that the observed synergy between CX-5461 and BMH-21 is driven by enhanced p53 activation. We hypothesize that the enhanced p53 activation by the combination treatment arises from dual activation via the nucleolar surveillance pathway (NSP) and DNA damage response (DDR) **(Figure 4G)**. Here, we found that the combination of CX-5461 and BMH-21 inhibits rRNA transcription more efficiently than either drug alone, indicating a potentiated effect on Pol I inhibition. These results demonstrate that CX-5461 and BMH-21 are mutually compatible in Pol I inhibition and suggests that these drugs are not functionally redundant. While the exact mechanisms of Pol I inhibition by each drug are not completely resolved, CX-5461 has been shown to inhibit Pol I transcription initiation, whereas BMH-21 has been shown to inhibit both initiation and elongation (Mars et al., 2020; Jacobs et al., 2022). This complementary inhibition likely contributes to their synergistic effects.

Strikingly, we observed that combining CX-5461 and BMH-21 reduces abundance of the Pol I large catalytic subunit RPA194. Previous studies have shown that BMH-21, unlike CX-5461, readily induces RPA194 proteasomal degradation (Peltonen et al., 2014; Pitts et al., 2022). We found that combining BMH-21 with CX-5461 promotes RPA194 downregulation compared to BMH-21 monotherapy, suggesting that CX-5461 might enhance RPA194 degradation by BMH-21. Additionally, CX-5461 downregulates the MYC oncogene, which transcriptionally regulates Pol I (Lee et al., 2017). Therefore, the downregulation of Pol I we observe with combination treatment may occur either during transcription or after translation. Regardless, the loss of RPA194 with combination treatment demonstrates that combining CX-5461 and BMH-21 enhances on-target inhibition of Pol I, leading to NSP and p53 activation. Interestingly, RPA194 expression has been recently linked to ferroptosis resistance by suppressing mitophagy-dependent iron release, and sensitization to ferroptosis by combined CX-5461 and BMH-21 treatment might contribute to the synergy we observe (Chang et al., 2025; Zhang et al., 2025). Given the abilities of CX-5461 and BMH-21 to interact with DNA, it would be interesting to assess in future studies whether these agents interact with mitochondrial DNA to sensitize cancer cells to ferroptosis or other non-apoptotic death pathways.

In addition to NSP activation downstream of Pol I inhibition, the DNA damage response (DDR) pathway might also contribute to p53 activation by combined CX-5461 and BMH-21 treatment. Indeed, we observed increased phosphorylation of H2A.X with combination treatment, consistent with enhanced induction of DNA damage. Simultaneous activation of p53 by converging NSP and DDR pathways may explain why p53 is significantly activated by combination treatment. However, increased DNA damage induction may also contribute to dose-limiting toxicity in patients. Therefore, future preclinical studies on Pol I inhibitor combinations are needed to evaluate adverse effects associated with DNA damage including hematological and gastrointestinal toxicities alongside occurrence of secondary malignancy (Martorana et al., 2022).

Our finding that CX-5461 and BMH-21 synergize in reducing cancer cell viability at low to moderate doses is clinically intriguing, as the synergy we observed in the combination enables significant dose reduction while maintaining efficacy. Even though the combination treatment was modestly synergistic (around 10-15% greater viability reduction), these combinations comprise substantially less of each drug than would be required to achieve a similar effect with either drug alone. This benefit is especially poignant considering that CX-5461 has been associated with significant patient adverse effects, including nausea and phototoxicity of the skin and eyes (Hilton et al., 2022). BMH-21 has not been assessed in human clinical trials, and therefore the toxicity of BMH-21 remains unknown. However, given that CX-5461 and BMH-21 are associated with different toxicity profiles, combined Pol I inhibition could provide clinical benefits compared to high-dose monotherapy, including better tolerability and reduction of dose-limiting toxicity. The potential for combination treatment to minimize adverse effects while maintaining efficacy makes this strategy particularly promising.

Combined RNA polymerase I inhibition with CX-5461 and BMH-21 represents a promising strategy for selectively targeting Pol I activity in cancer. Given that rRNA production is critical for ribosomal biogenesis, which is essential for the protein synthesis required by rapidly proliferating cancer cells, dual targeting Pol I represents a highly relevant approach in cancer therapy. Our findings that combined CX-5461 and BMH-21 treatment enhances on-target Pol I inhibition may extend to diseases other than cancer including aging and neurodegenerative disorders where Pol I inhibition has proved effective (LeDoux, 2024; Sharifi et al., 2024).

In our current study, the synergistic effect between CX-5461 and BMH-21 were observed when applying both drugs simultaneously. Therefore, we envision that these agents may be applied simultaneously in cancer treatment. In light of CX-5461’s ability to irreversibly inhibit Pol I initiation, it will be interesting to see in future studies if sequential therapy with CX-5461 and BMH-21 may also be effective (Mars et al., 2020). Although we limited our study solely on MCF-7 cells, it will be interesting in future studies to examine other cancer models to assess whether CX-5461 and BMH-21 exhibit synergy broadly across other breast cancer subtypes and cancer topologies. While future preclinical studies are required to evaluate the pharmacokinetics and dynamics of this combination, the observed synergy in reducing MCF-7 cancer cell viability provides strong rationale for continued development of this therapeutic strategy. With continued optimization and investigation, dual Pol I inhibition may offer a potent and broadly applicable treatment modality that is both effective and well-tolerated in the context of cancers and other diseases.

## 7 Conflict of Interest

The authors declare that the research was conducted in the absence of any commercial or financial relationships that could be construed as a potential conflict of interest.

## 8 Author Contributions

J.Y.C and B.A.K. conceived the project. J.Y.C. and K.N.N. designed and performed experiments. J.Y.C., K.N.N., and B.A.K analyzed the data. J.Y.C. wrote the initial draft and created the figures. K.N.N. and B.A.K. reviewed and edited the manuscript. B.A.K. supervised the project and secured funding.

## 9 Funding

The work was supported by grants from the U.S. National Institutes of Health NIGMS R01-GM141033 and Upstate Cancer Center Pilot Grant (B.A.K.).

## 10 Acknowledgments

We thank Lawrence I. Rothblum for his advice and experimental expertise regarding nascent RNA quantitation. We thank Ahmed M. Fakhr for his assistance with cell culture. We thank Ryan J. Palumbo and the members of the Knutson lab for their feedback and assistance in editing the work. We also thank the SUNY Upstate Medical University Department of Biochemistry and Molecular Biology and the SUNY Upstate MD/PhD program for their support.

## 12 Data Availability Statement

The original contributions presented in the study are included in the article/supplementary material, further inquiries can be directed to the corresponding author.

## Notes

### Competing Interest Statement

The authors have declared no competing interest.

