## Supplementary Material for "Dual RNA Polymerase I Inhibition with CX-5461 and BMH-21 Synergizes in Breast Cancer by Activating p53-Dependent Stress"

### 1 Supplementary Figures and Tables

#### 1.1 Supplementary Table

| CX-5461 Best-Fit IC50 | |
|---|---|
| BMH-21 (µM) | CX-5461 IC50 (µM) |
| 0 | 3.079 |
| 0.5 | 2.181 |
| 1 | 0.839 |
| 2 | 9.405 |
| 4 | 7.099 |

**Supplementary Table 1.** Low to moderate dose BMH-21 treatment enhances CX-5461 potency. CX-5461 best-fit IC50 values for cells treated with indicated doses of BMH-21. CX-5461 dose-response curves were fit using a 4-parameter logistic regression in GraphPad Prism. IC50 values reduced by BMH-21 treatment compared to cells treated with CX-5461 alone (increased CX-5461 potency) are colored red. IC50 values increased by BMH-21 treatment compared to cells treated with CX-5461 alone (reduced CX-5461 potency) are colored green.

### 1.2 Supplementary Figures

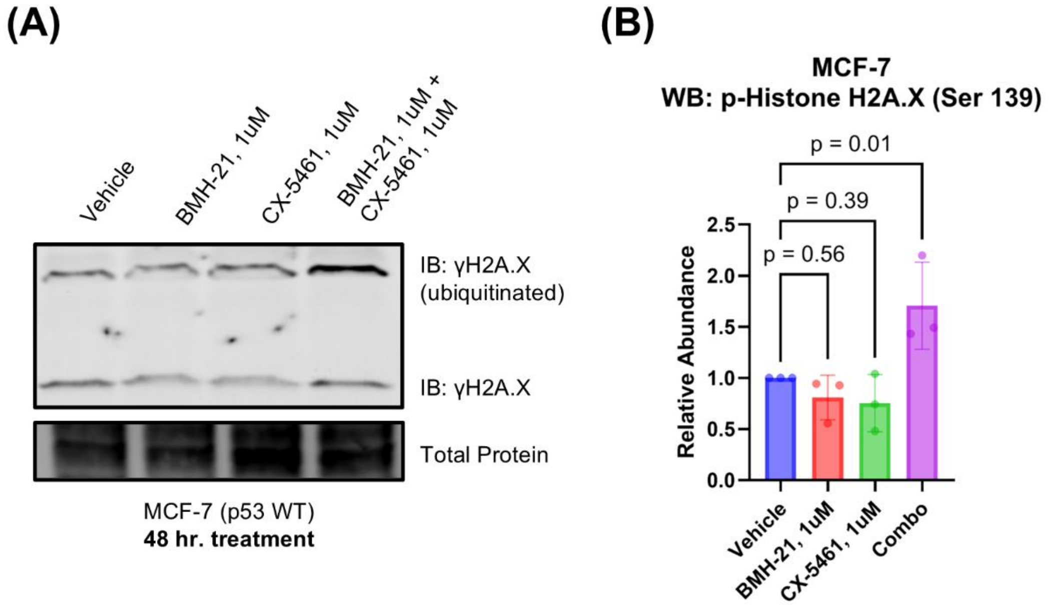


**Supplementary Figure 1.** CX-5461 and BMH-21 synergistic combinations enhance DNA damage. **(A)** Western blot probing for histone 2A family member X phosphorylation (γH2A.X) as a measure of DNA damage. MCF-7 cells were treated for 48 hours with elevated doses of CX-5461 and BMH-21 prior to harvesting. A representative Western blot is shown. **(B)** Densitometry of (A). Bars represent the average of three biological replicates and error bars represent the standard deviation. Relative abundance was compared using RM one-way ANOVA with Dunnett's multiple comparisons test.

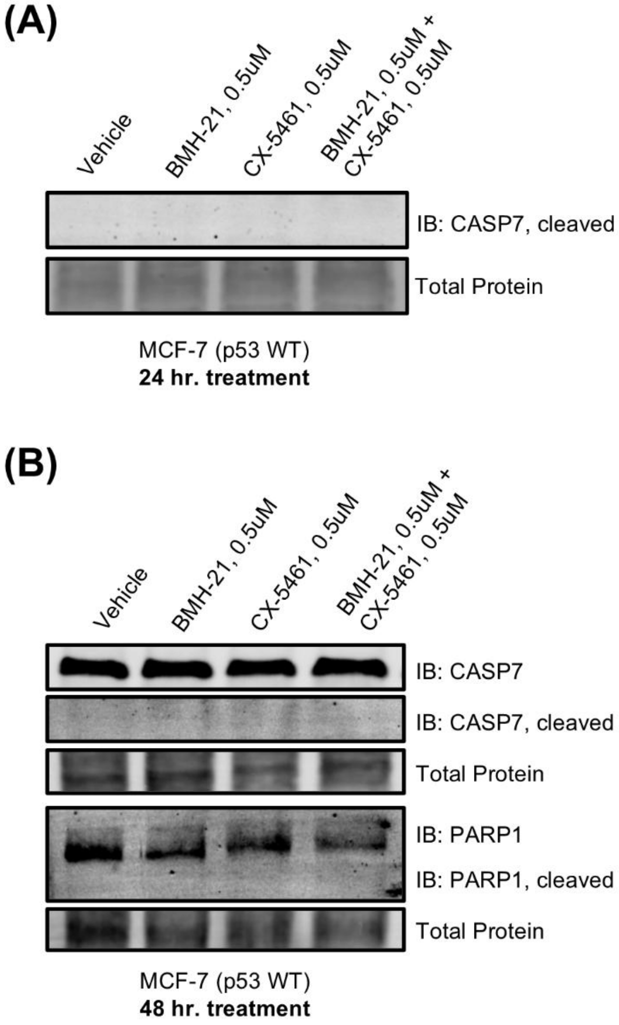


**Supplementary Figure 2.** Low synergistic doses of CX-5461 and BMH-21 do not induce detectable caspase-7 or PARP1 cleavage. (A) Western blot probing for cleaved caspase-7 (CASP7) as a measure of apoptosis induction. MCF-7 cells were treated for 24 hours with low synergistic doses of CX-5461 and BMH-21 prior to harvesting for Western blot. (B) Western blot probing for caspase-7 (CASP7) and poly(ADP-ribose) polymerase 1 (PARP1) total and cleaved forms as a measure of apoptosis induction. MCF-7 cells were treated for 48 hours with low synergistic doses of CX-5461 and BMH-21 prior to harvesting for Western blot.
